# The nuclear ubiquitin ligase adaptor SPOP is a conserved regulator of C9orf72 dipeptide toxicity

**DOI:** 10.1101/2021.03.09.434618

**Authors:** Carley Snoznik, Valentina Medvedeva, Jelena Mojsilovic-Petrovic, Paige Rudich, James Oosten, Robert G. Kalb, Todd Lamitina

## Abstract

A hexanucleotide repeat expansion in the C9orf72 gene is the most common cause of inherited amyotrophic lateral sclerosis (ALS) and frontotemporal dementia (FTD). Unconventional translation of the C9orf72 repeat produces dipeptide repeat proteins (DPRs). Previously, we showed that the DPRs (PR)50 and (GR)50 are highly toxic when expressed in *C. elegans* and this toxicity depends on nuclear localization of the DPR. In an unbiased genome-wide RNAi screen for suppressors of (PR)50 toxicity, we identified 12 genes that consistently suppressed either the developmental arrest and/or paralysis phenotype evoked by (PR)50 expression. All of these genes have vertebrate homologs and 7/12 contain predicted nuclear localization signals. One of these genes was *spop-1*, the *C. elegans* homolog of SPOP, a nuclear localized E3 ubiquitin ligase adaptor only found in metazoans. SPOP is also required for (GR)50 toxicity and functions in a genetic pathway that includes *cul-3*, which is the canonical E3 ligase partner for SPOP. Genetic or pharmacological inhibition of SPOP in mammalian primary spinal cord motor neurons suppressed DPR toxicity without affecting DPR expression levels. Finally, we find that genetic inhibition of *bet-1*, the *C. elegans* homolog of the known SPOP ubiquitination targets BRD2/3/4, suppresses the protective effect of SPOP mutations. Together, these data suggest a model in which SPOP promotes the DPR-dependent ubiquitination and degradation of BRD proteins. We speculate the pharmacological manipulation of this pathway, which is currently underway for multiple cancer subtypes, could also represent a novel entry point for therapeutic intervention to treat C9 FTD/ALS.

**Significance statement:** The G4C2 repeat expansion in the *C9orf72* gene is a major cause of Fronto-Temporal Dementia (FTD) and Amyotrophic Lateral Sclerosis (ALS). Unusual translation of the repeat sequence produces two highly toxic dipeptide repeat proteins, PR_X_ and GR_X_, which accumulate in the brain tissue of individuals with these diseases. Here, we show that PR and GR toxicity in both *C. elegans* and mammalian neurons depends on the E3 ubiquitin ligase adaptor SPOP. SPOP acts through the bromodomain protein BET-1 to mediate dipeptide toxicity. SPOP inhibitors, which are currently being developed to treat SPOP-dependent renal cancer, also protect neurons against DPR toxicity. Our findings identify a highly conserved and ‘druggable’ pathway that may represent a new strategy for treating these currently incurable diseases.

## Introduction

Mutations in *C9ORF72* are the most common monogenic cause of ALS and FTD (the most common non-Alzheimer’s adult dementia) (1, 2). The mutation is an expansion of the microsatellite repeat, 5’-GGGGCC-3’ in the intron between exons 1A and 1B. In unaffected individuals, the hexanucleotide repeat is reiterated 5 – 15 times, while in ALS/FTD patients, the hexanucleotide repeat expansion (HRE) occurs hundreds or even thousands of times. Three non-mutually exclusive C9 pathophysiological mechanisms have been proposed: 1) reduced abundance of the native C9 protein, 2) C9 pre-mRNA containing the HRE adopts an imperfect hairpin structure that sequesters RNA binding proteins, and 3) the generation of dipeptide repeat proteins (DPRs) by non-AUG-dependent translation (RANT) (3) of the HRE C9 pre-mRNA (4).

RANT of the sense G_4_C_2_ sequence, as well as a disease-associated antisense G_2_C_4_ sequence, leads to the production of five DPRs - Gly-Ala (GA), Gly-Pro (GP), Gly-Arg (GR), Pro-Ala (PA), and Pro-Arg (PR) (5). At autopsy, each of these DPRs is detected immunohistologically in C9 FTD and ALS patients but not non-C9 FTD or ALS patients. While DPR sites of expression and overall levels are highly variable, they are significantly more abundant within frontal and temporal lobe regions affected in FTD (6), suggesting a significant pathological role in this disease.

Of the five DPRs, two – Gly-Arg (GR) and Pro-Arg (PR) – are toxic in multiple cellular and animal model systems, although Gly-Ala (GA) can also be toxic in some, but not all, settings (7–9). Pathophysiological processes linked to DPRs include: 1) perturbation of nucleocytoplasmic shuttling (10, 11), 2) impaired protein translation (12), 3) Inhibition of proteasome function (13, 14), 4) defects in U2 snSNP-dependent splicing (15), 5) compromised mitochondrial function (16), and 6) perturbation of membraneless organelle dynamics (17, 18). In all of these studies, the key experimental approach involved the non-RANT expression of codon-varied DPRs, which allows experimental separation of G4C2 RNA toxicity from individual DPR toxicity. It is believed that these codon-varied DPR models accelerate the appearance of phenotypes that ordinarily emerge over decades in human C9 patients (7, 12, 17–25).

Here we describe an unbiased genetic screen for DPR modifiers in *C. elegans*. The genes identified in this screen included many previously identified DPR modifiers, as well as several novel genes. One novel (PR)50 suppressor was the E3 ubiquitin ligase adaptor **<u>S</u>**peckle-type Pox virus and Zinc Finger (**<u>PO</u>**Z) domain **<u>P</u>**rotein, or SPOP, which is a common genetic cause of prostate, endometrial, and renal cancer (26) but has never been linked to neurodegenerative disease. CRISPR-based introduction of SPOP missense alleles frequently found in prostate cancer also provided protection against (PR)50 toxicity. SPOP inhibition is a conserved mechanism to protect against DPR toxicity since either SPOP knockdown or an SPOP inhibitor in rat primary neurons also provided protection against DPR-induced neuron death. Finally, we find that the protective effects of SPOP inhibition depends on the bromodomain-containing protein and known SPOP substrate *bet-1*.

## Results

### A genome-wide RNAi screen for suppressors of DPR toxicity in *C. elegans* identifies known and novel conserved genes

Expression of C9orf72 DPRs (PR)50 and (GR)50 are toxic and cause cellular and organismal toxicity in multiple systems (7, 9, 18, 19, 21, 27). Expression of these DPRs in *C. elegans* muscles cells leads to a completely penetrant larval developmental arrest and paralysis phenotype (20). To identify genes required for this phenotype, we performed a genome-wide RNAi screen for suppressors of (PR)50 induced developmental arrest and/or paralysis. We screened 15,865 individual gene knockdowns and identified 391 initial (PR)50 suppressors. Each hit was rescreened in six independent trials. 12/391 hits showed suppression of the (PR)50 growth arrest in at least 4/6 of these trials (Table 1, Fig. S1). 83% of gene knockdowns (10/12) also suppressed toxicity caused by (GR)50 in at least 1/2 age-dependent paralysis assays (Table 1, Fig. S2). Since loss of these genes suppresses toxicity, these genes are necessary for DPR toxicity. The genes identified as (PR)50 suppressors fell into three broad functional classes; 1) Nuclear transport and RNA binding, 2) Ubiquitin-mediated protein degradation, and 3) chromatin regulation (Table 1). 50% (6/12) of these genes encode homologs of genes previously identified as modifiers of or interactors with (PR)50 or (GR)50 in yeast, fly, or mammalian cells (9, 18, 19, 21), suggesting (PR)50 engages similar pathophysiological mechanisms in *C. elegans* as it does in other systems. 75% (9/12) of (PR)50 suppressors are conserved from yeast to humans, while 25% (3/12) genes are absent from yeast but present in metazoan genomes. None of the genes are specific to *C. elegans*. Only one gene, *ergo-1*, appears to act via transgene suppression since knockdown reduces both GFP and RFP expression by >50% from a transgene expression control strain (Fig. S3), which is consistent with previous reports of *ergo-1(RNAi)* acting as a transgene suppressor (28). 75% of genes (9/12) contain either a predicted nuclear localization signal or experimental evidence for nuclear localization, suggesting the encoded protein is localized to the nucleus where (PR)50 is required for toxicity (Table 1) (20). 75% of genes (9/12) are annotated as having ‘lethal’ and/or ‘sterile’ phenotypes, suggesting that traditional forward genetic screening for DPR suppressors would not have isolated these genes since these screens usually require mutants to be viable and fertile in order to be identified (Table 1). Therefore, our RNAi screen identified highly conserved and essential genes required for (PR)50 toxicity, many of which are likely to be localized to the nucleus, where (PR)50 is also required for toxicity.

**Table 1.** RNAi gene knockdowns that suppress myo-3p::(PR)50-GFP toxicity.

| *C. elegans* | Yeast | Fly | Human | Description | Predicted or demonstrated NLS | Suppresses (GR)50? | Lethal/Sterile |
| --- | --- | --- | --- | --- | --- | --- | --- |
| **Nuclear transport / RNA binding** |  |  |  |  |  |  |  |
| *rsp-6* | MRD1 | X16<sup>2</sup> | SRSF3<sup>1</sup> | RNA binding protein | Yes | Yes | Yes |
| *ztf-4* | NGR1 | CG42458<sup>2</sup> | NCOA5/hnRNPC<sup>1</sup> | contains RRM domain, similar to hnRNPC | Yes | Yes | Yes |
| *gyf-1* | SYH1 | Gigyf | GIGYF<sup>1</sup> | yeast homolog SYH1 binds proline-rich sequences and influences nuclear pore distribution | Yes | Yes | No |
| *ima-3* | KAP122<sup>4</sup> | Kap-α3<sup>2,3</sup> | KPNA3<sup>1</sup> | 1 of 3 nuclear importin alpha subunits | No | No | Yes |
| *ergo-1* | None | AGO1T1 | AGO3 | Production of endo-siRNAs | Yes | Yes | No |
| **Ubiquitin-mediated protein degradation** |  |  |  |  |  |  |  |
| *spop-1* | None | roadkill | SPOP | Speckle-type POZ protein; no yeast homolog | Yes | Yes | No |
| *ufd-2* | Ufd2 | UBE4B/CG9934 | UBE4a | E3 and/or E4 ubiquitin ligase | Yes | Yes | No |
| *rpn-12* | RPN12 | RPN12 | PSMD8<sup>5</sup> | nonATPase regulatory subunit of the 26S proteasome | No | Yes | No |
| *rpn-9* | RPN9 | RPN9 | PSMD13 | nonATPase regulatory subunit of the 26S proteasome | No | Yes | Yes |
| **Chromatin regulation** |  |  |  |  |  |  |  |
| *met-2* | MET2 | eggless | SETDB1 | Histone methyltransferase | Yes | Yes | Yes |
| *utx-1* | CYC8 | Utx | UTX/UTY<sup>6</sup> | Histone demethylase | Yes | Yes | Yes |
| *pis-1* | None | PTIP | PAXIP1 | Pax transcription factor interacting protein | Yes | No | Yes |

1 – Interacts with both PR and GR in human cells (18)
2 – Enhanced the toxicity of (GR)50 in *Drosophila* model (18)
3 – Enhanced the toxicity of (PR)25 in *Drosophila* model (21)
4 – Suppresses the toxicity of (PR)50 when overexpressed in yeast (9)
5 – Gene is enriched or depleted in (PR)20 treated K562 cell CRISPR screen (19)
6 – Gene is enriched or depleted in (GR)20 treated K562 cell CRISPR screen (19)

We obtained previously characterized mutant alleles that were viable and fertile for several of the genes identified in our screen. To validate our RNAi results, we crossed two of these mutants, *met-2(n4256)* and *rsp-6(ok798)*, into the (PR)50 background. As we found for RNAi knockdown, both loss-of-function mutants suppressed the developmental arrest phenotype and the post-developmental age-dependent paralysis phenotype caused by (PR)50 (Fig. 1C). We examined (PR)50 localization in each of these mutants and found that nuclear localization of the DPR was disrupted. In either mutant background, (PR)50-GFP was distributed throughout the non-nuclear compartments and was largely absent from either the nucleus or nucleolus (Fig. 1D). These data provide independent validation of the hits derived from our RNAi screen and suggest that the mechanism of suppression for some genes is associated with re-localization of (PR)50 out of the nucleus.

**Figure 1.**
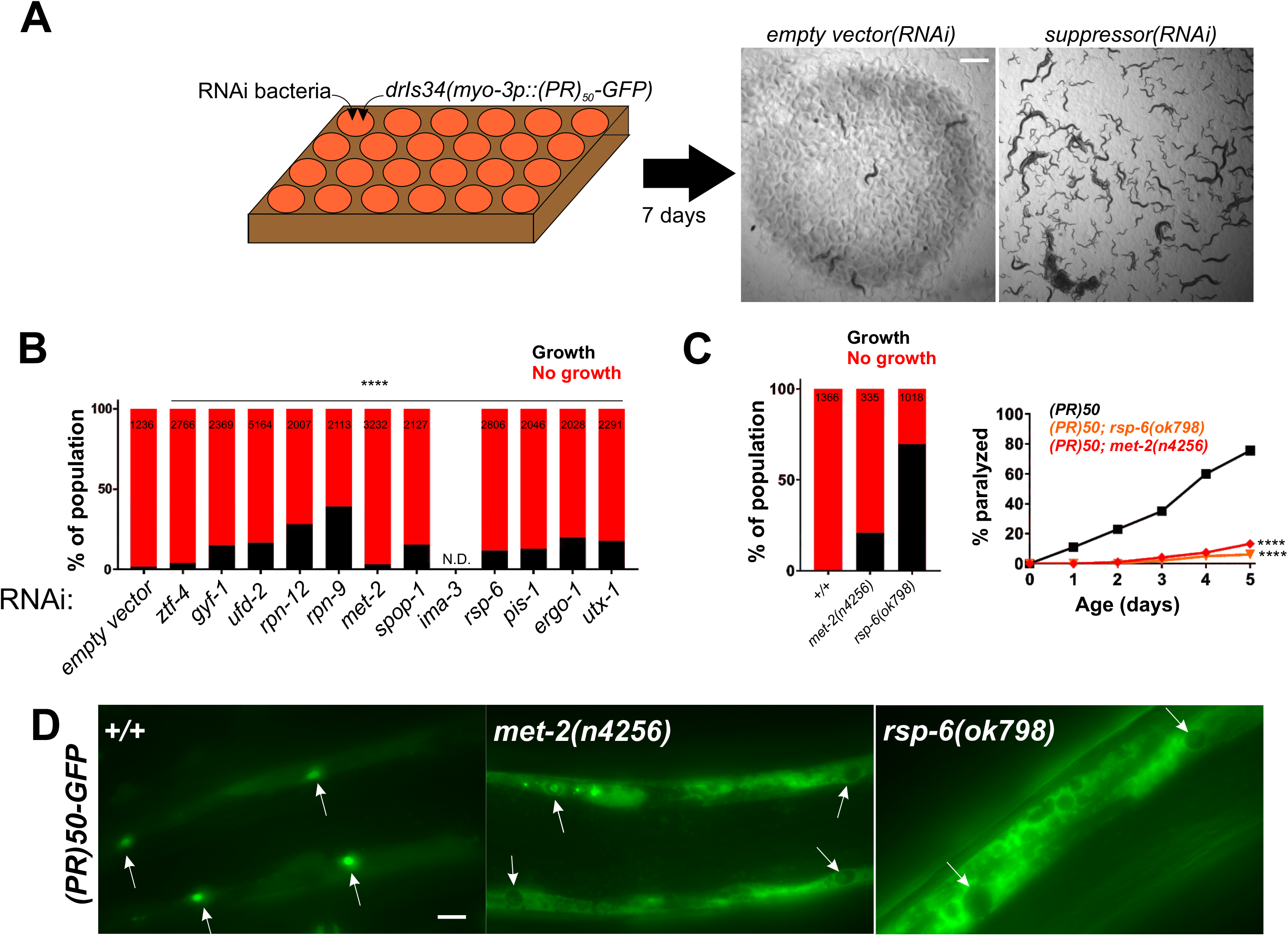
A genome-wide RNAi screen for suppressors of (PR)50 toxicity in *C. elegans.* A) RNAi screening strategy. ∼30 hypochlorite isolated eggs from the *drIs34 (myo-3p::(PR)50-GFP; myo-3p::mCherry*) strain are seeded onto 24 well RNAi plates. After 7 days, each well was visually examined and scored relative to the *empty vector(RNAi)* control, which exhibits no growth and strong paralysis. Suppressors are identified as wells in which worms continue to grow and/or regain motility (*gfp(RNAi*) shown as an example). B) COPAS quantification of worm length (time-of-flight(TOF)) in RNAi suppressors of (PR)50. Worms with a TOF >200 were placed in the ‘Growth’ group while those with TOF <200 were placed in the ‘No growth’ group. Data are presented as the percentage of the total population in either the ‘Growth’ or ‘No growth’ groups. The size of each population (N) is indicated at the top of each bar. **** - p<0.0001, Fisher’s exact test versus *empty vector(RNAi)*. N.D. – not determined due to completely penetrant lethal phenotype in this replicate. C) Age-dependent paralysis assay was performed with homozygous viable mutations in genes identified in the RNAi screen. N>50 for each genotype. **** - p<0.0001, Log-Rank test with Bonferroni correction. D) Localization of (PR)50-GFP in either the WT or indicated homozygous mutant background. Arrows point to muscle nucleus. In WT, (PR)50-GFP is primarily localized to nuclei but in *n4256* and *ok798*, (PR)50-GFP is largely excluded from the nucleus and accumulates in non-nuclear compartments. Scale bar = 10 microns.

### Loss-of-function mutations in the E3 ubiquitin ligase adaptor SPOP suppress (PR)50 and (GR)50 toxicity

In addition to several previously identified DPR modulators (*rsp-*6/SRSF3, *ima-3/*KPNA3, *ztf-4*/hnRNPC, *gyf-1*/GIGYF, *rpn-12*/PSMD8, *utx-1*/UTX/UTY), our screen also identified several new genes associated with regulation of ubiquitin proteasome activity. One newly identified gene was *spop-1*, the *C. elegans* homolog of human speckled-type POZ domain containing protein (SPOP) (Fig. 2A,B). *spop-1(RNAi)* robustly suppressed the age-dependent toxicity of both (PR)50 and (GR)50 (Fig. 2C,D). We obtained or generated via CRISPR/Cas9 mutations in *spop-1* (*gk630214, dr28*) and crossed them into (PR)50 and (GR)50 expressing *C. elegans*. *spop-1(gk630214)* is a nonsense mutation early in the *spop-1* coding sequence that truncates the protein within the MATH domain and is therefore likely a null allele (Fig. 2A). *spop-1(dr28)* is a CRISPR engineered 82bp deletion within exon 2 that deletes the start codon for isoform b and creates a reading-frame shift in the isoform a, also creating a likely null allele (Fig. 2A). Both *spop-1* mutations recapitulated the RNAi phenotypes observed for both age-dependent paralysis (Fig. 2E,F) and growth arrest (Fig. 2B). To determine if *spop-1* also functions to regulate neuronal DPR toxicity, we utilized a newly developed line expressing (GR)50 in the motor neurons. Motor neuron (GR)50 caused a progressive, age-dependent loss in motility (Fig. S4). The *spop-1(gk630214)* mutant suppressed this motor neuron (GR)50 phenotype (Fig. S4). Therefore, loss of *spop-1* function suppresses C9orf72-associated DPR toxicity in multiple cellular contexts.

**Figure 2.**
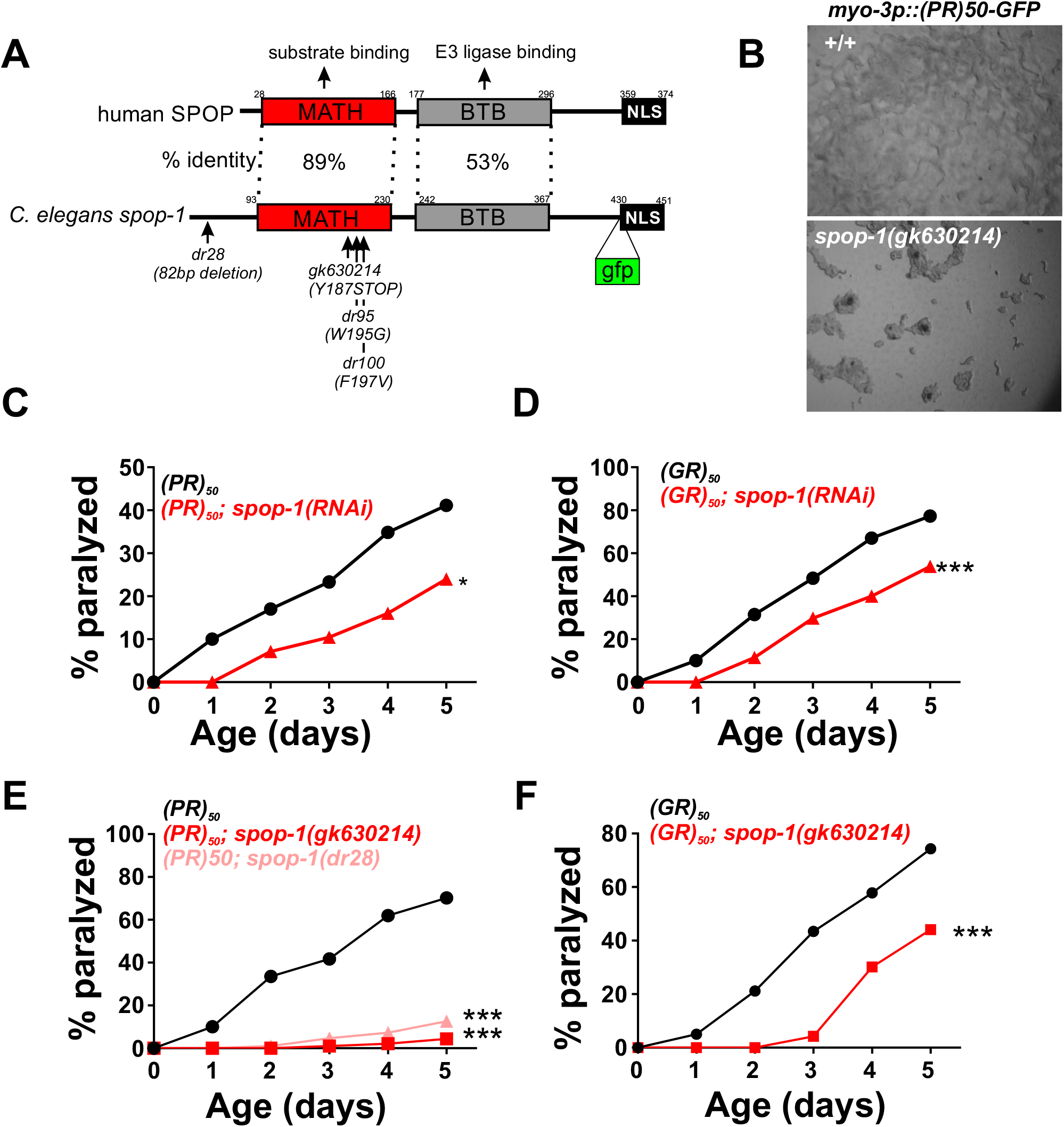
Loss of function mutations in the E3 ubiquitin ligase adaptor *spop-1* suppress (PR)50 and (GR)50 toxicity in *C. elegans*. A) Domain structure of the SPOP protein. The percent identity between the human and *C. elegans* domains are shown. The *dr28* allele is a CRISPR-induced allele that deletes 82bp within the second exon of *spop-1*. This shifts the reading frame of isoform 1 to encounter a premature stop codon and deletes that start ATG for isoform 2. Therefore, *dr28* is a likely *spop-1* null allele. The *gk630214* allele is a *spop-1* nonsense mutation isolated from the Million Mutation project. The *gfp* insertion site used to make the SPOP-1-GFP CRISPR allele is shown. B) *spop-1(gk630214)* and *ufd-2(tm1380)* suppress the developmental arrest phenotype of (PR)50-GFP. C,D) *spop-1(RNAi)* suppresses the age-dependent paralysis phenotype of (PR)50-GFP (C) an (GR)50-GFP (D). E,F) *spop-1* alleles *gk630214* and *dr28* suppress (PR)50-GFP (E) and (GR)50-GFP (F). For all paralysis assays, N=50-100 animals for each genotype, * - p<0.05, *** - p<0.001, Log-rank test with Bonferroni correction for multiple testing.

In mammals, SPOP functions as an adaptor protein that targets substrates for ubiquitination by cullin 3-type E3 ubiquitin ligases and subsequent proteasomal degradation (26). If *spop-1* confers protection against DPR toxicity through its canonical function as an E3 ubiquitin ligase adaptor, then inhibition of the cullin 3 ubiquitin ligase should phenocopy *spop-1* mutants and protect against DPR toxicity. Furthermore, this protective effective should exhibit a non-additive gene interaction with *spop-1* mutants. Since *cul-3* mutants are not viable, we tested these predictions by performing *cul-3(RNAi)* in the wild type and *spop-1(dr28)* mutant background. *cul-3(RNAi)* in a wild type background provided significant protection against (PR)50 toxicity (Fig. 3A). While *spop-1(dr28)* also suppressed (PR)50 toxicity, *cul-3(RNAi)* in the *spop-1(dr28)* mutant did not provide additional suppression (Fig. 3A). This is unlikely to be due to a floor effect since *gfp(RNAi)* consistently suppresses (PR)50 toxicity to a greater extent than *spop-1(dr28)* (Fig. 3A). Together, these genetic data suggest that *cul-3* functions in the same genetic pathway as *spop-1* to mediate DPR toxicity in *C. elegans*.

**Figure 3.**
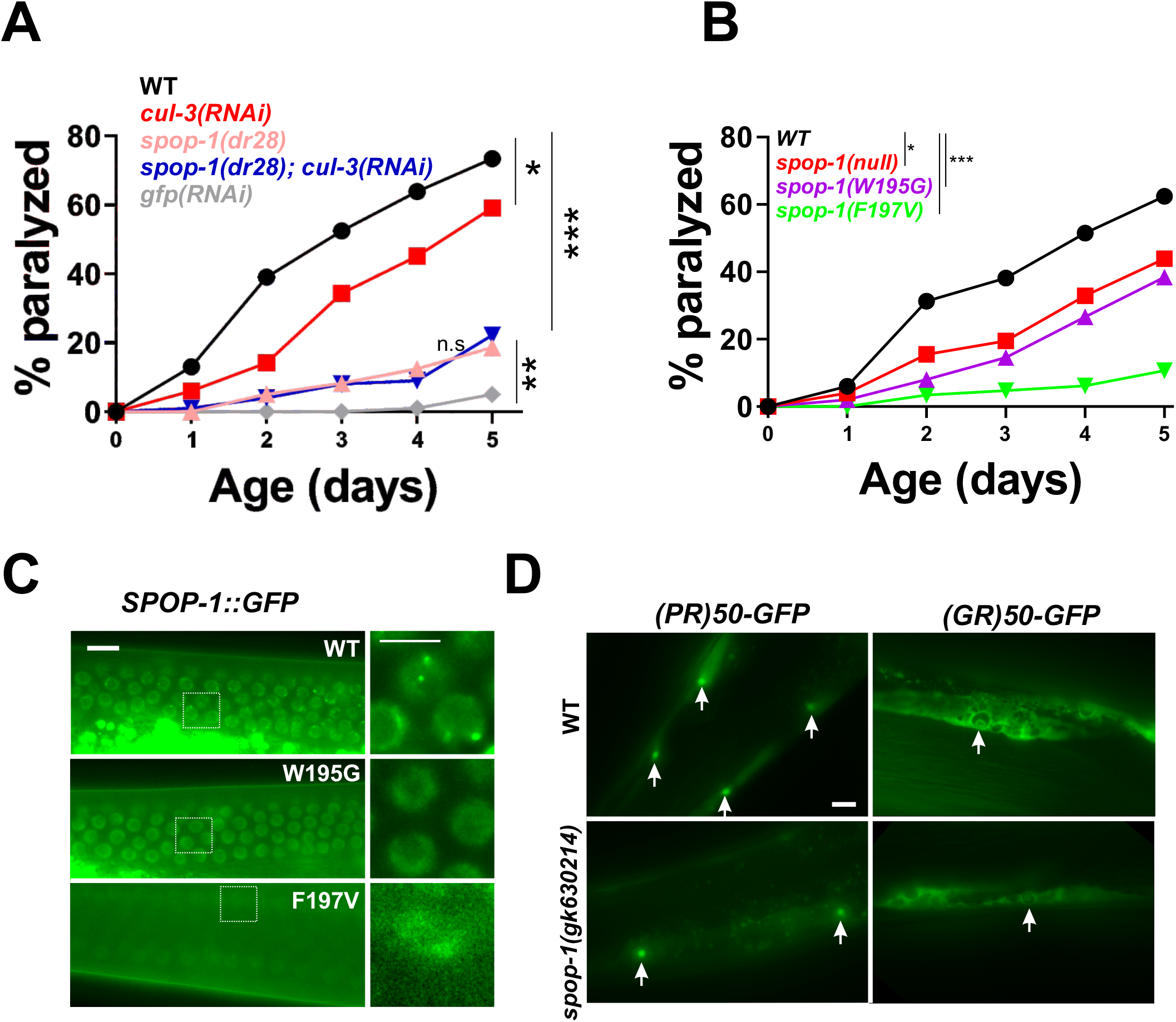
Inhibition of a *spop-1-cul-3* pathway protects against DPR toxicity without altering DPR levels or localization. A) (PR)50 paralysis assay in the indicated genetic backgrounds. N=100 animals per genotype. *-p<0.05, **- p<0.01, ***-p<0.001, 1 way ANOVA with Tukey post-hoc comparisons. B) (PR)50 paralysis assay in the indicated *spop-1* genotypes. Mutations were introduced into the endogenous *C. elegans spop-1* gene via CRISPR/Cas9. N=100 animals per genotype. *-p<0.05, ***-p<0.001, 1 way ANOVA with Tukey post-hoc comparisons. C) Localization of a CRISPR-engineered SPOP-1-GFP fusion protein containing the indicated point mutations. Germline nuclei are shown because their high density facilitates observation of multiple instances of SPOP-1-GFP speckling. Similar speckling behavior is observed in somatic cells. Speckling was not observed in SPOP-1-GFP^W195G^. SPOP-1-GFP^F197V^ expression was at the limit of detection and speckling could not be evaluated. Scale bar = 10 microns. Boxed regions are enlarged, deconvolved images to illustrate SPOP-GFP speckling. Scale bar in enlarged image = 5 microns. D) (PR)50-GFP and (GR)50-GFP expression in wild type or *spop-1(gk630214)*. Images are exposure matched. Arrows point to muscle nuclei with nucleolar DPR enrichment, as was previously described for (PR)50-GFP and (GR)50-GFP localization in *C. elegans* (20). Scale bar = 10 microns.

### Cancer-causing missense mutations in SPOP protect against (PR)50 toxicity

Genetic missense mutations in SPOP are a common cause of prostate cancer. The most common mutations, W131G and F133V, are loss-of-function. These mutations are in the SPOP MATH domain and interfere with the ability of SPOP to interact with its substrates. If the role of SPOP in DPR toxicity is to target substrates for *cul-3*-dependent ubiquitination and degradation, then introduction of these mutations into *C. elegans spop-1* is predicted to phenocopy the *spop-1* null alleles and confer protection against (PR)50 toxicity. We used CRISPR/Cas9 to introduce each mutation into the endogenous *spop-1* gene (W195G and F197V in *C. elegans*) in the (PR)50 background. We also knocked these mutations into a *spop-1-gfp* allele tagged at the endogenous locus with CRISPR/Cas9. Like the *spop-1* null alleles, both missense mutations conferred protection against (PR)50 toxicity (Fig. 3B). While *spop-1^W195G^-gfp* exhibited normal abundance and nuclear localization, *spop-1^F197V^-gfp* showed a substantial decrease in the SPOP-1-GFP protein levels, suggesting the F197V mutation reduces SPOP-1 protein levels (Fig. 3C). Loss of function *spop-1* alleles did not alter the levels or localization of either the (PR)50 or (GR)50 proteins, suggesting *spop-1* functions downstream of DPR nuclear localization to mediate toxicity (Fig. 3D). Together, these genetic data are consistent with the hypothesis that *spop-1* mediates DPR toxicity via *cul-3*-dependent ubiquitination and degradation of one or more *spop-1* substrates.

### SPOP is required for DPR toxicity in mammalian neurons

SPOP is highly conserved from *C. elegans* to humans (Fig. 1), but is absent from yeast. To determine if the role of SPOP in DPR toxicity is evolutionarily conserved, we utilized a rat spinal cord neuron culture system model of DPR toxicity. Embryonic rat spinal cord motor neurons were infected with recombinant Herpes Simplex Virus (HSV) engineered to express miRNAs (miRs) targeting rat SPOP. Two independently derived SPOP miRs led to a ∼90% reduction in SPOP protein levels in pure neuronal cultures (Fig. 4A). After growing neurons on an astrocyte feeder layer for two weeks, we then co-infected our spinal cord neuronal cultures with two separate HSVs expressing a codon-optimized DPRs (GA50, (GR)50, or (PR)50) and the SPOP miR. In prior work, we show that both viruses express their cargo in >95% of neurons in a non-toxic manner (29, 30). We found that expression of known toxic DPRs GA50, (GR)50, and (PR)50 caused significant neuronal toxicity and reduced the number of viable primary motor neurons ∼50% (Fig. 4B). Expression of a scrambled control sequence miR was neither toxic nor protected against the toxicity of (PR)50 (Fig. 4B). However, expression of the SPOP miR provided highly significant neuroprotection against the toxicity of all DPRs. Interestingly, while the SPOP miR robustly inhibited SPOP protein expression, the levels of several known SPOP ubiquitination substrates identified in cancer cells (e.g., PTEN, DUSP7, and G3BP1) were not altered by either (PR)50, (GR)50, or SPOP knockdown (Fig. 4C). Since these SPOP substrates have been largely characterized in renal, prostate, and endometrial cancer cell lines, these data present the possibility that neuronal SPOP functions may include the degradation of novel neuronal substrates. Finally, as we saw in *C. elegans spop-1* mutants, SPOP knockdown in mammalian neurons did not alter the abundance of any of the toxic DPRs (either nominal monomers or higher molecular weight species; Fig. 4D). These data suggest that SPOP is an evolutionarily conserved factor broadly required for DPR toxicity from *C. elegans* to mammalian neurons and that it functions downstream of DPRs to engage mechanisms leading to cellular toxicity.

**Figure 4.**
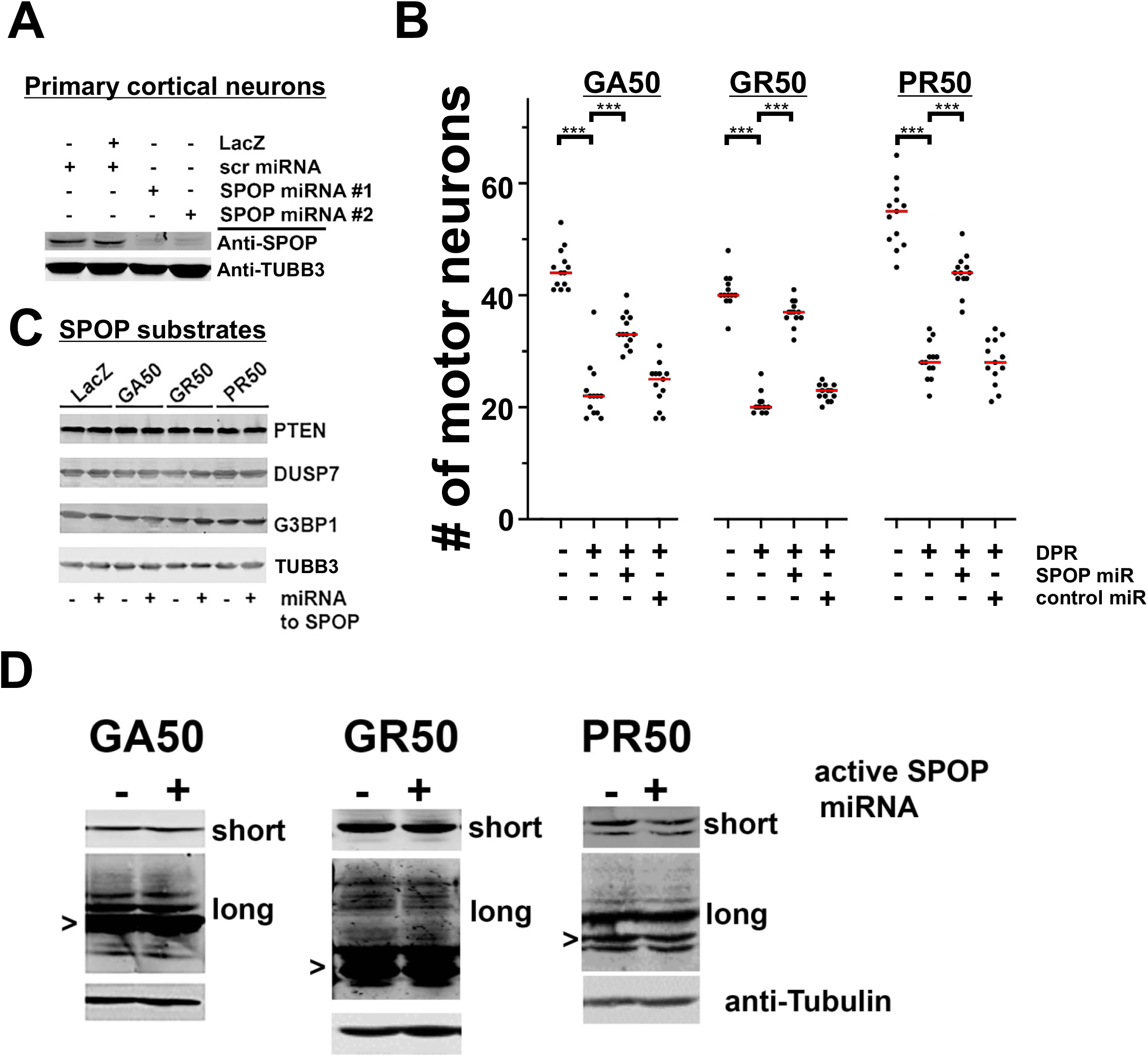
SPOP knockdown protects against DPR toxicity in mammalian primary neurons without reducing the abundance of known SPOP ubiquitination targets or DPRs themselves. A) Cortical neuron cultures were infected with Herpes Simplex Virus (HSV) engineered to express LacZ, a scrambled sequence microRNA (miR) or two different miRs targeting SPOP. Endogenous neuronal SPOP levels are reduced by targeted miRNAs. B) Days *in vitro*14 (DIV14) spinal cord neuron cultures were co-infected with HSV engineered to express the indicated DPR and either the scrambled sequence miR (control – “cont”) or miR targeting SPOP. Control cultures were infected with HSV-LacZ. Five days later, cultures were processed for immunocytochemistry and motor neuron counts were obtained. By ANOVA, DPRs lead to a statistically significant reduction in motor neuron number (compared with LacZ expressing cultures) and knockdown of SPOP leads to a statistically significant protection against SPR toxicity (e.g., GA(50) group, *F*(*_3,48_*) = 84.8, p < 0.001; GR(50) group, *F*(*_3,47_*) = 240.6, p < 0.0001; PR(50) group, *F*(*_3,48_*) = 119.0, P < 0.0001). C) Cortical neuron cultures were infected with HSV engineered to express LacZ or the GA(50), (PR)50 or (GR)50 DPRs. Simultaneously, cultures were infected with either the HSV engineered to express the scrambled sequence miR or SPOP miR. Two days later, before any neuron loss is detected, lysates were probed for known SPOP targets. No statistically significant group differences were detected in scan band intensities by ANOVA (not shown). D) Lysates from experiments described in C) were probed for GA, GR or PR and imaged with short or long exposures (to see less abundant higher molecular weight species). Putative monomers are noted with “>”. No statistically significant group differences were detected in scan band intensities by ANOVA (not shown).

SPOP is strongly upregulated in most clear cell renal carcinomas and genetic inhibition of SPOP in these cancer cells attenuates their growth (31). As a result, Guo et al performed a small molecule screen for SPOP inhibitors (32). One inhibitor, Compound 6b, blocks SPOP-substrate interactions with low micromolar affinity. We asked whether 6b was neuroprotective in our tissue culture model. Application of 6b had no effect on motor neuron survival in naive cultures (not shown). However, 6b exhibited a statistically significant blunting of DPR toxicity in cultures that express (GR)50, GA50 or the exogenously provided PR20 peptide (Fig. S5) (14, 19, 33). These observations suggest that the SPOP-CUL3 pathway we have defined may be amenable to small molecule modulation in mammals.

### The bromodomain containing protein BET-1 is required for *spop-1* mutant suppression of DPR toxicity

Our data suggest a model in which a conserved function of SPOP leads to the ubiquitin-dependent degradation of one or more substrates when toxic DPRs are present. In an SPOP mutant, this substrate(s) is no longer ubiquitinated and degraded, leading to increased protein levels and protection against DPR toxicity. A prediction of this model is that reducing the abundance of this substrate should reverse the protective effect of *spop-1* mutants against DPRs. To identify such substrates, we performed a *spop-1* mutant suppressor screen. In mammalian cancer cells, *spop-1* targets >30 proteins for degradation (26). Eleven of these substrates have homologs in *C. elegans*. We performed feeding-based RNAi and paralysis assays for 9 of these candidate SPOP substrates in both the WT and *spop-1(dr28)* mutant. Only a single gene knockdown against the BRD2/3/4 homolog *bet-1* suppressed *spop-1(dr28)* protection against DPR toxicity (Fig. 5). In wild type animals, *bet-1(RNAi)* did not further enhance DPR toxicity (Fig. 5A). *bet-1(RNAi)* also suppressed the paralysis phenotypes of the *spop-1* LOF allele *gk630214* (Fig. 5B) and the *spop-1* W195G missense allele (Fig. 5C). This indicates that the *bet-1(RNAi)* suppression of *spop-1* is not allele specific. Rather, loss of *bet-1* suppresses the phenotypic consequences associated with loss of *spop-1*. Taken together, these genetic data are consistent with a model in *spop-1* mediates the degradation of *bet-1* and that stabilization of *bet-1* protects against DPR toxicity (Fig. 5D).

**Figure 5.**
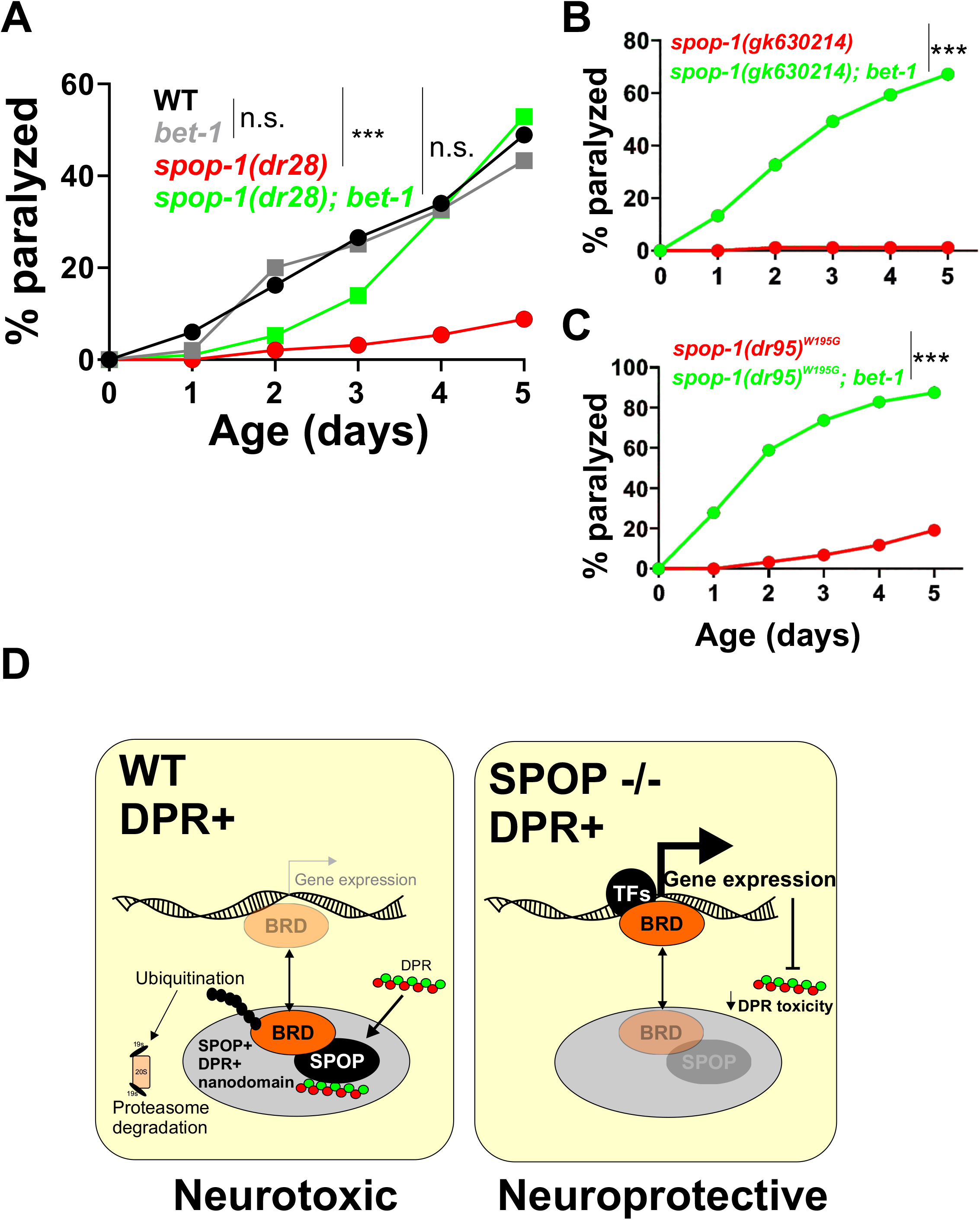
*spop-1* mutant protection against DPR toxicity requires the bromodomain protein *bet-1*. A) Paralysis assay with the indicated genotypes. *bet-1* null alleles are homozygous sterile (59). Therefore, *bet-1(RNAi)* was utilized. All genotypes include the *drIs34 (myo-3p::3XFLAG-(PR)50-GFP)* transgene. N=100 animals per genotype. ***-p<0.001, n.s. – p>0.05, Log-Rank test with Bonferroni correction for multiple comparisons. B) Paralysis assay in the indicated *spop-1* allele +/− *bet-1(RNAi)*. N=100 animals per genotype. ***-p<0.001, Log-Rank test with Bonferroni correction for multiple comparisons C) Paralysis assay in the indicated *spop-1* allele +/− *bet-1(RNAi)*. N=100 animals per genotype. ***- p<0.001, Log-Rank test with Bonferroni correction for multiple comparisons D) Model for the function of SPOP and BRD proteins in C9 DPR toxicity.

## Discussion

Aggregated, misfolded, and otherwise insoluble proteins, including DPRs, accumulate in the brains of individuals who bear the HRE in the C9ORF72 locus and these species are widely believed to be key contributors to disease pathophysiology. One potential driver of this process is direct inhibition of the proteasome by DPRs. Cryo-electron tomographic imaging of cells expressing GA DPRs show that twisted GA filaments decorated with proteasomes, which appear arrested in a non-functional state (13). Similarly, PR DPRs directly bind to purified proteasomes and inhibit degradation of ubiquitinated substrates (14). Here we define components of a novel pathway that also ties the ubiquitin-proteasomal degradation pathway to DPR toxicity. In our unbiased RNAi screen, we discovered that the E3 ubiquitin ligase adaptor SPOP and its associated E3 ligase, cullin 3, are required for DPR toxicity. The beneficial effects of ablation of SPOP, as well as introduction of mutations that inactivate its ability to present clients to the E3 ligase, provide strong evidence that the canonical ubiquitination function of SPOP is indispensable for DPR toxicity.

Previously, we have shown that the toxicity of PR and GR DPRs are dependent upon nuclear localization (20). This aligns well with the fact that SPOP is also a predominantly nuclear protein and suggests that SPOP-dependent degradation of a nuclear protein(s) renders cells vulnerable to DPR toxicity. SPOP was originally identified in 1997 as a POZ domain containing protein that localizes to subnuclear speckles (34). Subsequent studies revealed that SPOP speckles have biophysical characteristics associated with proteins that undergo liquid-liquid phase separation (LLPS) (35). Moreover, SPOP nuclear speckles are highly dynamic and are enriched with known SPOP ubiquitination substrates (36). Given that PR and GR dipeptides can both undergo LLPS themselves and interfere with LLPS associated with other proteins (17, 18, 37), our discovery that SPOP is required for DPR toxicity suggests the possibility that alterations in the LLPS properties of SPOP speckles by (PR)50 could be the basis for DPR toxicity. These alterations could either occur through direct or indirect SPOP-DPR interactions. For example, DPRs could directly interact with SPOP and block its ability to co-localize with and degrade specific ubiquitination targets. An alternative hypothesis is that DPRs could drive SPOP to inappropriately interact with and degrade targets that would not normally be degraded. Our data demonstrating that inhibition of the SPOP substrate *bet-1* suppresses the protective effect of SPOP inhibition on DPR toxicity supports the latter hypothesis. Additional studies examining SPOP-DPR interactions, as well as the roles of other SPOP substrates in mammalian cells, are needed to further differentiate between these possibilities.

Our studies are the first to link SPOP to a neurodegenerative disease. However, SPOP has well characterized links to non-neuronal diseases, such as prostate, endometrial, and renal cancer (31, 38, 39). In prostate cancer, recurrent somatic missense mutations such as W131G and F133V occur in ∼10% of metastatic patients (40). These mutations lead to SPOP loss of function and stabilization of multiple SPOP substrates (41, 42). Many additional missense mutations are associated with SPOP-dependent cancer, including several gain-of-function mutations associated with endometrial cancer that enhance the degradation of SPOP substrates (43). A prediction of our findings is that such gain-of-function alleles may enhance DPR toxicity. The highly conserved nature of the SPOP protein from *C. elegans* to humans, combined with efficient CRISPR engineering in worms and highly penetrant DPR phenotypic assays, provide a unique opportunity to define SPOP structure-function relationships in SPOP-dependent DPR toxicity in ways that are not possible in other systems.

SPOP targets many proteins for degradation (44). In SPOP prostate cancer mutants, the ability to target and degrade these substrates is lost, which leads to their upregulation (36). Upregulation of several such SPOP substrates are associated with different aspects of the cancer phenotype (43, 45, 46). One important set of substrates are the Bromodomain and Extra-Terminal motif (BET) containing proteins BRD2, BRD3, and BRD4. BET proteins are transcriptional co-activators that interact with acetylated histones and transcription factors to control gene expression (47). Our studies here show that BET proteins are also likely to play a role in SPOP-dependent DPR toxicity. We discovered that the protective effect of *spop-1* mutants on DPR toxicity was suppressed by reducing the abundance of the *C. elegans* protein *bet-1*. Our findings suggest a model in which DPRs lead to the inappropriate *spop-1*-dependent degradation of *bet-1*, and possibly other substrates. Loss of *spop-1* leads to BET-1 upregulation, which protects against DPR toxicity via alterations in gene expression. Inhibition of *bet-1* in the *spop-1* mutant suppresses these compensatory transcriptional responses and restores DPR toxicity. While our genetic studies are consistent with this model, additional cell biological and biochemical tests are needed to more rigorously test its predictions. Interestingly, in a recent small molecule screen, two BET inhibitors were identified as suppressors of exogenous PR20 toxicity (48). These observations are nominally the opposite effect our model would predict. The disparate results may be due to the differences in discovery platforms, read-out measures, redundant effects on multiple BRD domain proteins, and/or off-target drug effects. In this respect, genetic studies may be a more precise tool for investigating the role of BET proteins in DPR toxicity.

The primary goal of research into disease mechanism is to find a druggable target. In this regard, SPOP is an attractive candidate. The SPOP inhibitor Compound 6b stops the growth of SPOP overexpressing ccRCC cells in culture and attenuates tumor growth in a mouse model. A recent study identified 6b derivatives that are even more potent (49), suggesting that the SPOP pathway is highly suitable for small molecule intervention. Given that genetic inhibition of the cancer-relevant functions of SPOP can also suppress DPR toxicity, we predict that 6b and its derivatives could also be a new and potentially viable therapeutic option for opposing DPR toxicity. Preclinical studies show that 6b is well tolerated and has no major side effects (32), although the blood-brain barrier permeability of this compound is not known. Future studies investigating 6b or other SPOP inhibitors could provide a novel entry point for new ALS/FTD therapies.

## Conclusion

In conclusion, our unbiased genetic screening approach in *C. elegans* discovered that the SPOP ubiquitin ligase pathway is essential for C9 DPR toxicity. Given that SPOP is a nuclear protein and DPR nuclear localization is required for toxicity, our findings provide additional evidence that the mechanism of DPR toxicity originates in the nucleus. Drugs targeting both SPOP and its relevant targets, such as BET domain proteins, are useful tools for treating cancer and could be equally useful as C9 disease therapeutics.

## Materials and methods

### *C. elegans* strains, culture, and genome-wide RNAi screen

See Table 2 for a list of the strains utilized in this study. Unless otherwise indicated, animals were cultured at 20°C. The genome wide RNAi screen utilized the Ahringer feeding library (Source Bioscience, Nottingham, U.K.) and was performed as previously described (50, 51). Briefly, 96 well plates containing ∼100µl of LB + 25 µg/ml carbenicillin liquid media were seeded with individual bacterial clones using a 96 well pin tool and grown for 18-24 hours at 37°C. 20µl of this overnight culture was seeded into a single well of a 24 well plate containing NGM agar + 25 µg/ml carbenicillin + 1mM IPTG (Fisher Scientific) and incubated overnight at room temperature (∼20°C). Prior to seeding on RNAi screening plates, the *drIs34* strain is fed *gfp(RNAi)* bacteria to suppress the expression of (PR)50-GFP, which allows drIs34 animals to grow and reproduce. When fed *empty vector(RNAi)* or standard OP50, the *drIs34* strain produces progeny that are growth arrested and paralyzed. This phenotype is 100% penetrant. Therefore, *gfp(RNAi)* serves as a positive control for the (PR)50-GFP suppression screen and *EV(RNAi)* serves as a negative control. ∼30 synchronized L1 stage *drIs34* worms were seeded into each well and incubated at 20°C. After 7 days, each well was manually screened for RNAi clones that allowed further growth and/or restored mobility compared to the *EV(RNAi)* negative control. ‘Hits’ from this primary screened were re-screened six times and clones that were identified as ‘hits’ in 4/6 rescreens were considered validated hits. During the screening process, the experimenters were blinded to the identity of the RNAi clone. RNAi clones from validated hits were sequenced to determine the identity of the gene targeted. These sequence-validated strains were frozen as glycerol stocks and used for all subsequent RNAi-based experiments.

**Table 2.** Strains used in this study.

| **Strain name** | **Genotype** | **Reference** |
| --- | --- | --- |
| **OG736** | *drls28 [myo-3p::3XFLAG-(GR)50-gfp; myo-3p:dsRed2]* | (20) |
| **OG755** | *drls34 [myo-3p::3XFLAG-(PR)50-gfp; myo-3p:dsRed2]* | (20) |
| **OG919** | *met-2(n4256); drls34* | This study |
| **OG922** | *rsp-6(ok798); drls34* | This study |
| **VC40422** | *spop-1(gk630214)* | (61) |
| **OG1053** | *spop-1(gk630214) drls34* | This study |
| **OG1054** | *spop-1(gk630214); drls28* | This study |
| **OG1055** | *spop-1(dr26(CRISPR inserted gfp))* | This study |
| **OG1057** | *spop-1(dr28); drls34* | This study |
| **OG1153** | *drls49 [unc-47p::3XFLAG-(GR)50-gfp; unc-47p::gfp; myo-3p::mCherry]* | This study |
| **OG1158** | *spop-1(dr95(W195G)); drls34* | This study |
| **OG1159** | *spop-1(dr26 dr96(W195G))* | This study |
| **OG1168** | *spop-1(dr100(F197V); drls34* | This study |
| **OG1169** | *spop-1(dr26 dr101(F197V))* | This study |
| **OG1171** | *spop-1(gk630214); drls49* | This study |

### Molecular Biology and CRISPR methods

All procedures involving recombinant or synthetic nucleic acid molecules and materials were approved by the University of Pittsburgh Institutional Biosafety Committee. The *drIs34* ((PR)50) and *drIs28* ((GR)50) transgenes were previously described (20). CRISPR/Cas9 engineering was performed according to the method of Ghanta et al (52). For point mutations and deletion edits, worms were injected with a mixture of Cas9 (250ng/µl (IDT)), tracrRNA (100 ng/µl (IDT)), guide RNA (56 ng/µl (IDT)), repair oligonucleotide (110 ng/µl (IDT)), and *rol-6(su1006)* marker plasmid (40 ng/µl). The Cas9, tracrRNA, and guide RNA was incubated at 37°C for 10 minutes prior to the addition of the repair oligo and *rol-6* plasmid. For generation of the *spop-1-GFP* CRISPR allele, a gfp PCR product was PCR amplified from pPD95.75 (Addgene) using primers containing 35bp of overlap with the *spop-1* gene immediately upstream of the predicted nuclear localization sequence. Guide RNAs, repair oligonucleotides, PCR primers, and genotyping primers used in this study are listed in Table 3.

**Table 3.**
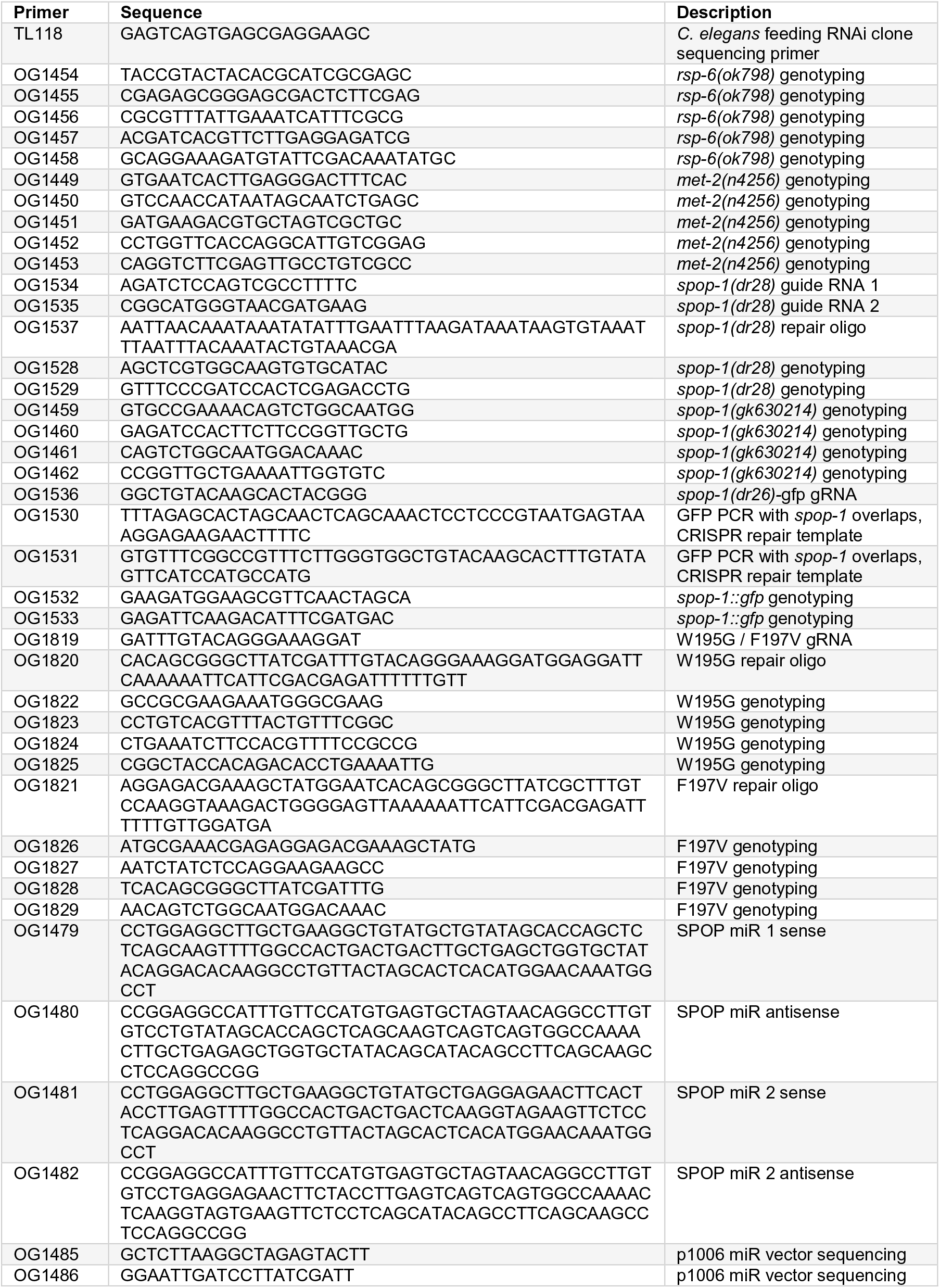
Oligonucleotides used in this study.

### Microscopy

Low magnification images were captured on a MZ16FA fluorescence stereo microscope equipped with a DFC345 FX camera and AF imaging software (Leica Microsystems). For high magnification images, day 1 adult hermaphrodites were anesthetized in 10 mM levamisole and imaged within a silicon grease enclosed slide chamber. Animals were imaged on a wide-field DMI4000B inverted microscope with a 63X oil immersion lens. Images were captured with a DFC 340 FX digital camera using AF imaging software (Leica Microsystems). All images were captured with identical exposure times and image brightness/contrast/intensity was adjusted for all images within a panel simultaneously within CorelDraw X7.

### Paralysis and thrashing assays

DPR paralysis assays were performed as previously described (20, 53). For the thrashing assay quantification of the integrated (GR)50 motor neuron line, 10 L4 animals were picked onto 3cm NGM plates spotted with OP50 (5 plates per genotype). At each time point, animals were picked into a dish containing ∼1ml of M9 solution. After a 5’ acclimation to liquid, the number of thrashes in 30 seconds was quantified. Each time the nose of the animals crossed the body midline was considered a thrash. 10 worms per genotype were measured at each time point. At the completion of the assay, the thrashing rates for each genotype at each time point were normalized to the ‘Day 0’ thrashing rate for that genotype in order to control for any baseline differences in thrashing. The experimenter counting the thrash rates was blinded to the genotype. Each experiment was performed 2-3 times and the results from a single representative experiment are shown.

### Mammalian neuronal cultures

Primary neurons cultures were established according to previously described methods (14, 29, 30). Briefly, cortical astrocytes from newborn rat pups were plated on German Glass, acid washed glass coverslips (18 millimeter round), grown in Minimal Essential Media MEM) + 10% fetal bovine serum and 10% horse serum (HS) and when they achieved ∼ 60% confluency, dissociated embryonic (E) day 15 rat spinal cord neurons were added. The media was changed to astrocyte-conditioned MEM + 10% HS + glial derived neurotrophic factor, ciliary neurotrophic factor and cardiotropin-1 (all at 2 nM). AraC (10uM) was then added to arrest astrocyte proliferation and washed out with subsequent bi-weekly media supplementation. After 14 days in vitro (DIV), recombinant HSV were added and cultures were fixed five days later. The SPOP miR clone was designed using the BLOCK-IT miR RNAi design tool (SPOP accession number NM_001100496.1; ThermoFisher Scientific). Sense and antisense oligos for three distinct rat SPOP miRs beginning at SPOP cDNA basepair 212, 239, and 336 were synthesized and cloned into the miR expression vector p1006 for packaging into HSV. Western blot validation experiments in HEK cells using overexpressed SPOP showed that the 336-miR was slightly less effective than the 212- or 239-miR. Therefore, only the 212-miR and 239-miRs were utilized. Viruses were generated by the Gene Delivery Technology Core at Massachusetts General Hospital.

For pure neuron biochemical studies, E17 rat cortex was dissociated and plated on plastic petri dishes and maintained in astrocyte-conditioned, Neurobasal + B27 supplement. After 14 DIV recombinant HSV was added and lysates prepared for western blotting. Antibodies used were: anti-GA (1:1000; Millipore Sigma), anti-GR (kind gift of Q. Zhu, Cleveland lab), anti-PR (kind gift of Y. Zhang, Petrucelli lab), anti-SPOP (1:500; Proteintech), anti-tubulin (1:5000; BioLegend), anti-PTEN (1:1000; Cell Signaling Technology), anti-DUSP7 (1:1000; Abcepta), and anti-G3BP1 (1:1000; Cell Signaling Technology).

#### Motor neuron toxicity assay

Methodological details of the in vitro mammalian motor neuron toxicity assay have been reported previously (29, 30, 54, 55). In brief, coverslips containing paraformaldehyde fixed astrocyte/spinal cord co-cultures were stained with SMI32 antibody – when restricted to neurons with a cell body diameter of 20 microns or greater, a validated marker of alpha motor neurons in this culture system (56). The number of motor neurons in three randomly selected 20x lower power field was determined and averaged. The average value of four coverslips was determined (experimental replicates). Determination of motor neuron number was obtained with the counter blinded to the experimental manipulation. The results from 3+ independent experiments are presented.

### Statistical Analysis

Paralysis assays were analyzed using the Kaplan-Meier log-rank function (OASIS (57, 58)). Comparisons of means were analyzed with either a two-tailed Students t-test (2 groups) or 1- or 2-way ANOVA (3 or more groups) using the Tukey or Dunn’s post-test analysis as indicated in GraphPad Prism 7 (GraphPad Software, Inc., La Jolla, CA). p-values of <0.05 were considered significant.

## Acknowledgements

This work was supported by grants from the NIH (NS094921 and NS096319 to T.L., NS05225 and NS087077 to R.G.K). Research in the Kalb lab is support by the Les Turner ALS Center. We thank the lab of Dr. Caiguang Yang (Shaghai Institute of Materia Medica, Chinese Academy of Sciences) for generously providing Compound 6b, the Petrucelli lab (Mayo Clinic) for generously providing the anti-GA antibody, and the Cleveland lab (UCSD) for generously providing the anti-GR antibody. Some strains were provided by the CGC, which is funded by the NIH Office of Research Infrastructure Programs (P40 OD010440).

---

**Figure S1.**
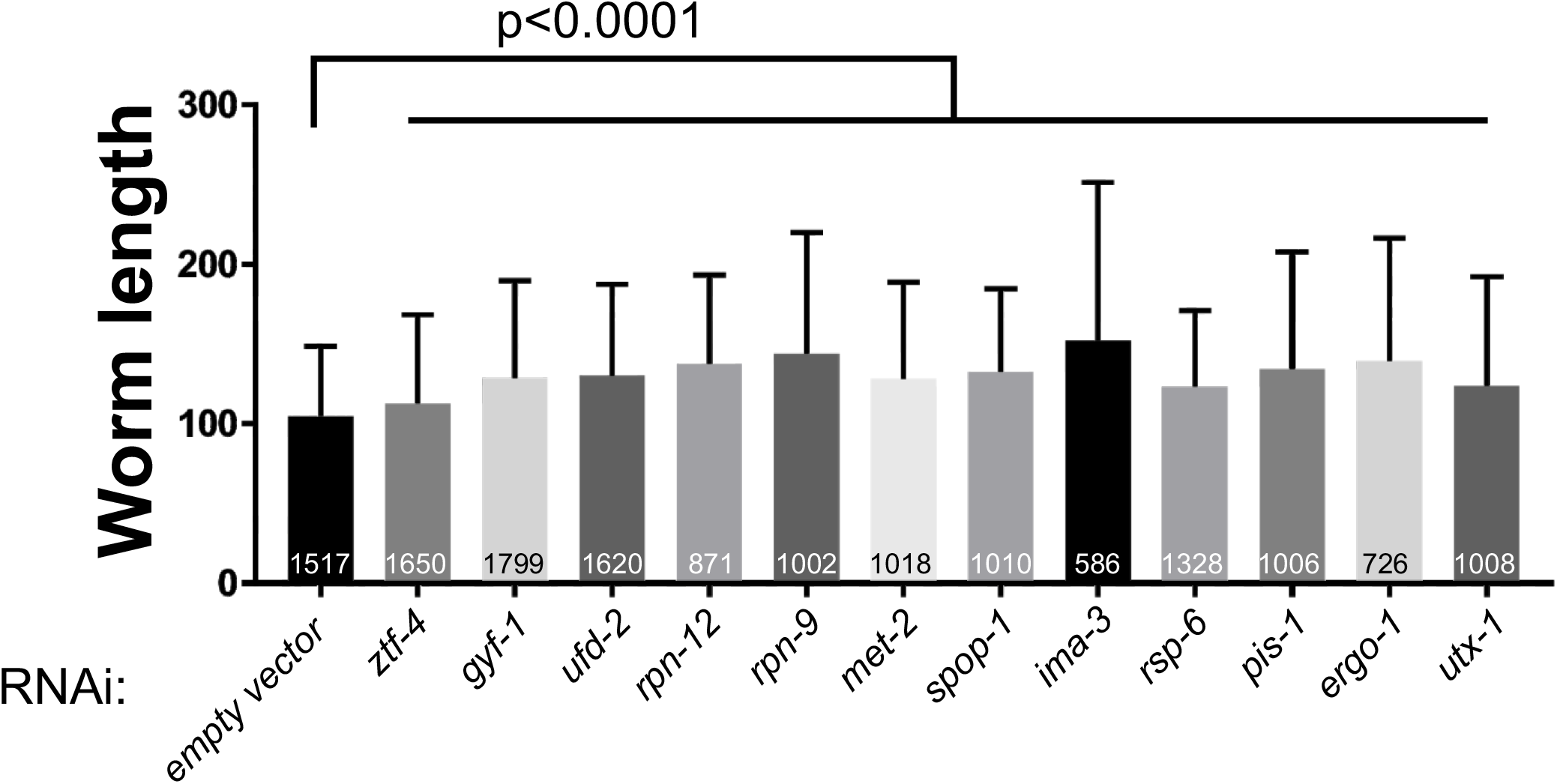
Quantification of suppression of (PR)50-induced growth arrest by hits from the genome-wide RNAi screen. *drIs34* eggs were placed on either (empty vector) RNAi or the indicated gene-specific RNAi bacteria. After 7 days at 20°C, animals were washed off the plate and worm length was quantified as the event time-of-flight using a COPAS Biosort. Data shown are mean +/− S.D. The N for each sample is shown on each bar. P-values versus the empty vector control were calculated using a non-parametric one-way ANOVA with Dunn’s multiple comparison post-hoc testing.

**Figure S2.**
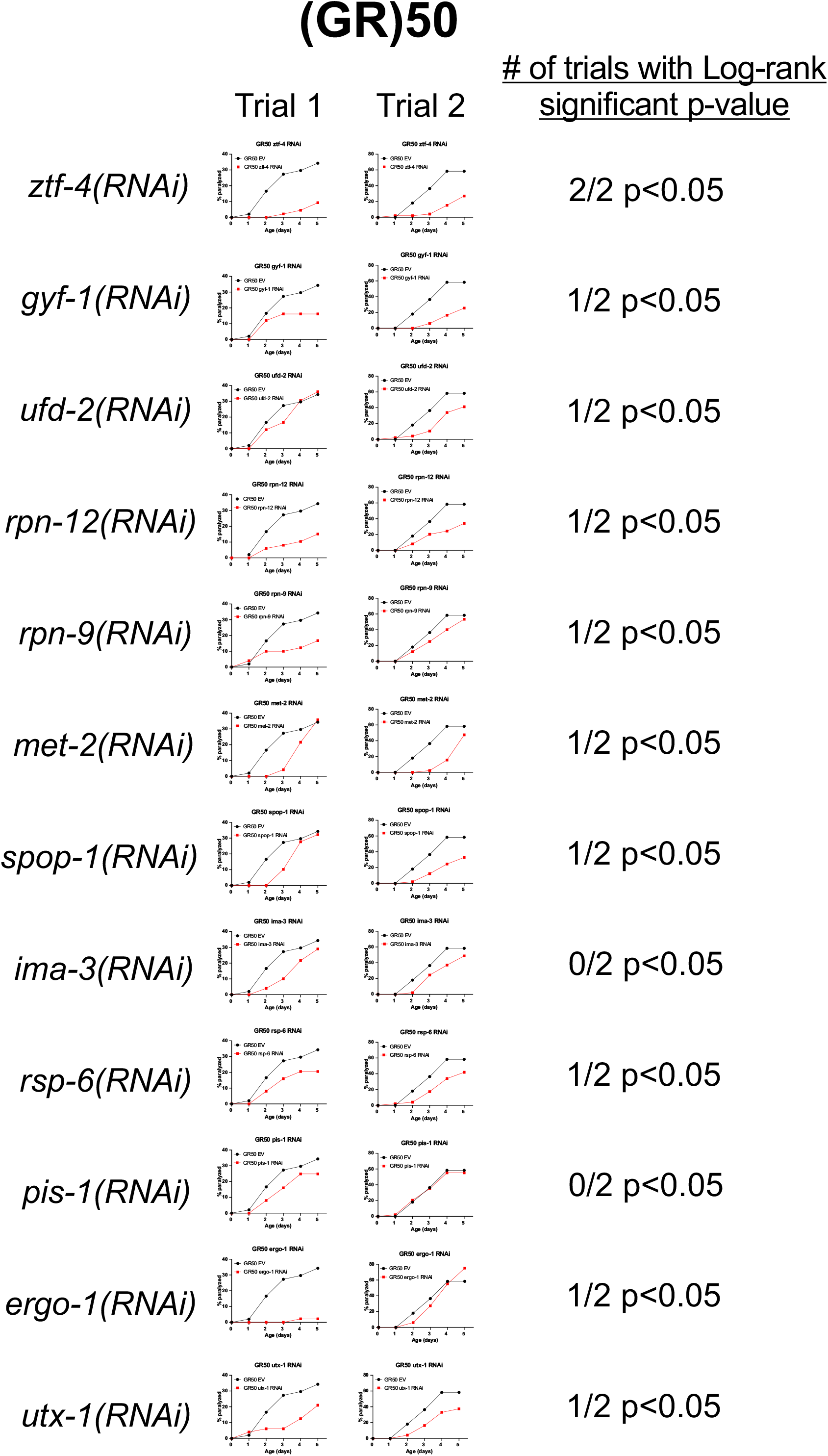
Effect of (PR)50 RNAi screen hits on (GR)50 age-dependent paralysis phenotype. Paralysis assays were performed as previously described (60). N=50-80 animals per assay. *EV(RNAi)* control in black and gene specific RNAi in red. Each candidate gene was scored in two separate assays and the number of times the Log-rank Bonferroni-adjusted p-value was <0.05 is indicated.

**Figure S3.**
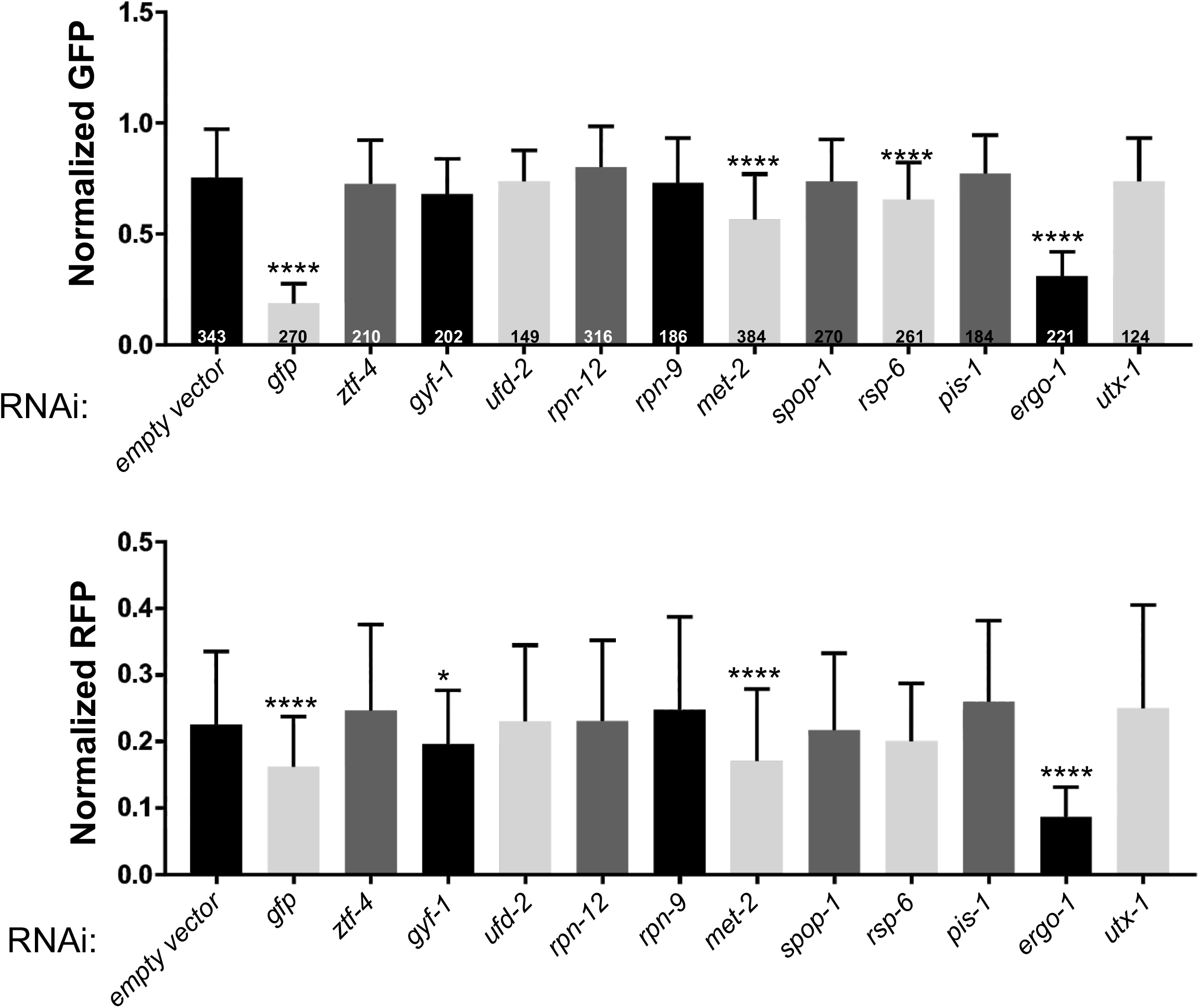
Effect of RNAi screen hits on transgene expression. *drIs33* eggs were placed on either *empty vector(RNAi)* or the indicated gene-specific RNAi bacteria. After 7 days at 20°C, animals were washed off the plate and normalized GFP (GFP/TOF; top) and normalized RFP (RFP/TOF; bottom) from adult animals with TOF>400 were quantified with a COPAS Biosort. Data shown are mean +/− S.D. * - p<0.05, **** - p<0.001, non-parametric One-way ANOVA with Dunn’s multiple comparisons test. The N for each sample is indicated within each bar.

**Figure S4.**
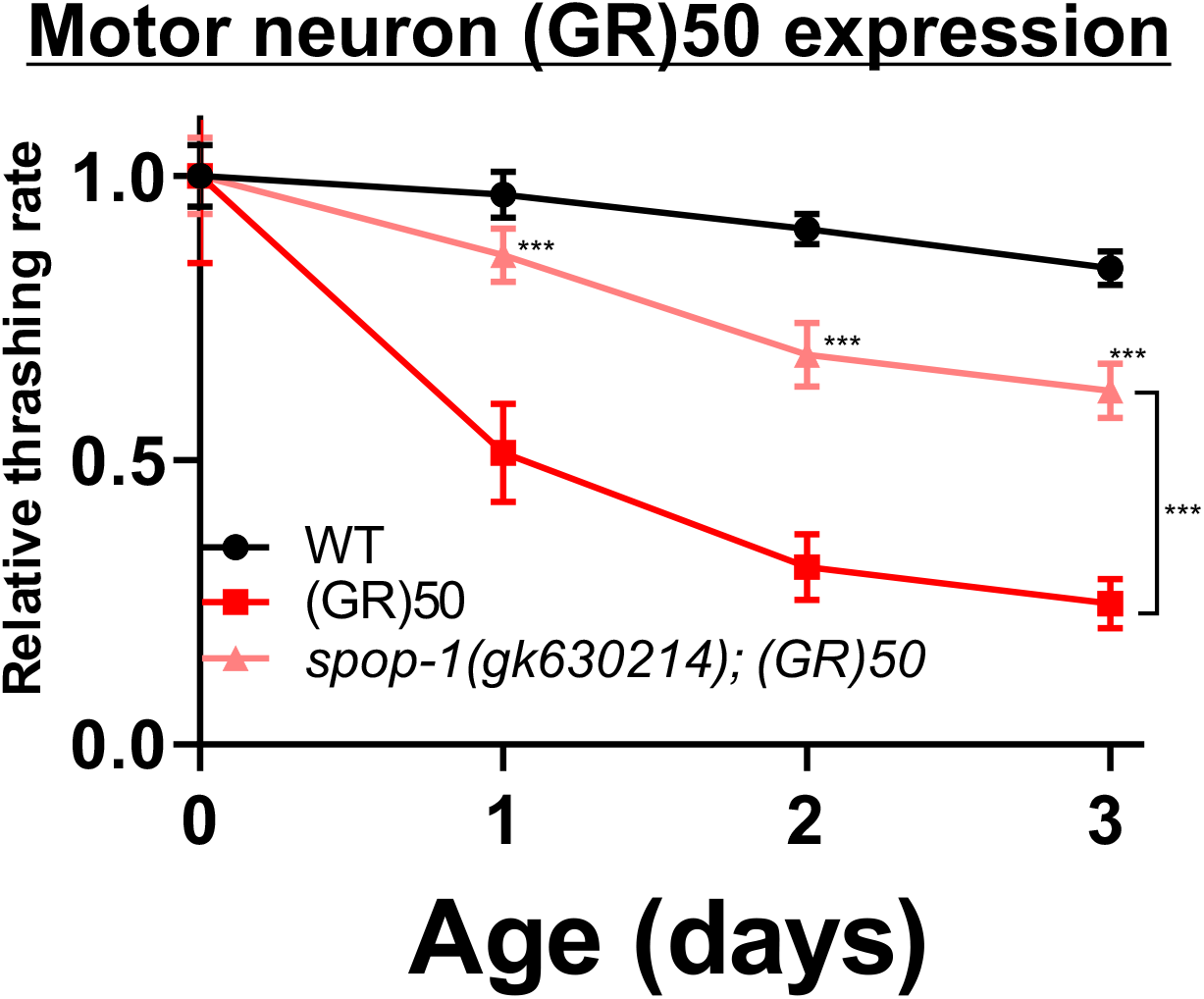
*spop-1* functions in motor neurons to protect against motor neuron expressed (GR)50-GFP. WT or *drIs49* (unc-47p::3xFLAG-(GR)50-GFP) animals of the indicated genotype were grown at 25°C and L4 stage animals (Day 0) were picked to new plates (4 sets of plates with 10 animals each per genotype). At each timepoint, the number of thrashes were quantified as previously described (60). The experimenter was blinded to genotype. Data within genotypes were normalized to the thrashing rate at Day 0. Data shown are the mean ± S.D., N=10. ***-p<0.001.

**Figure S5.**
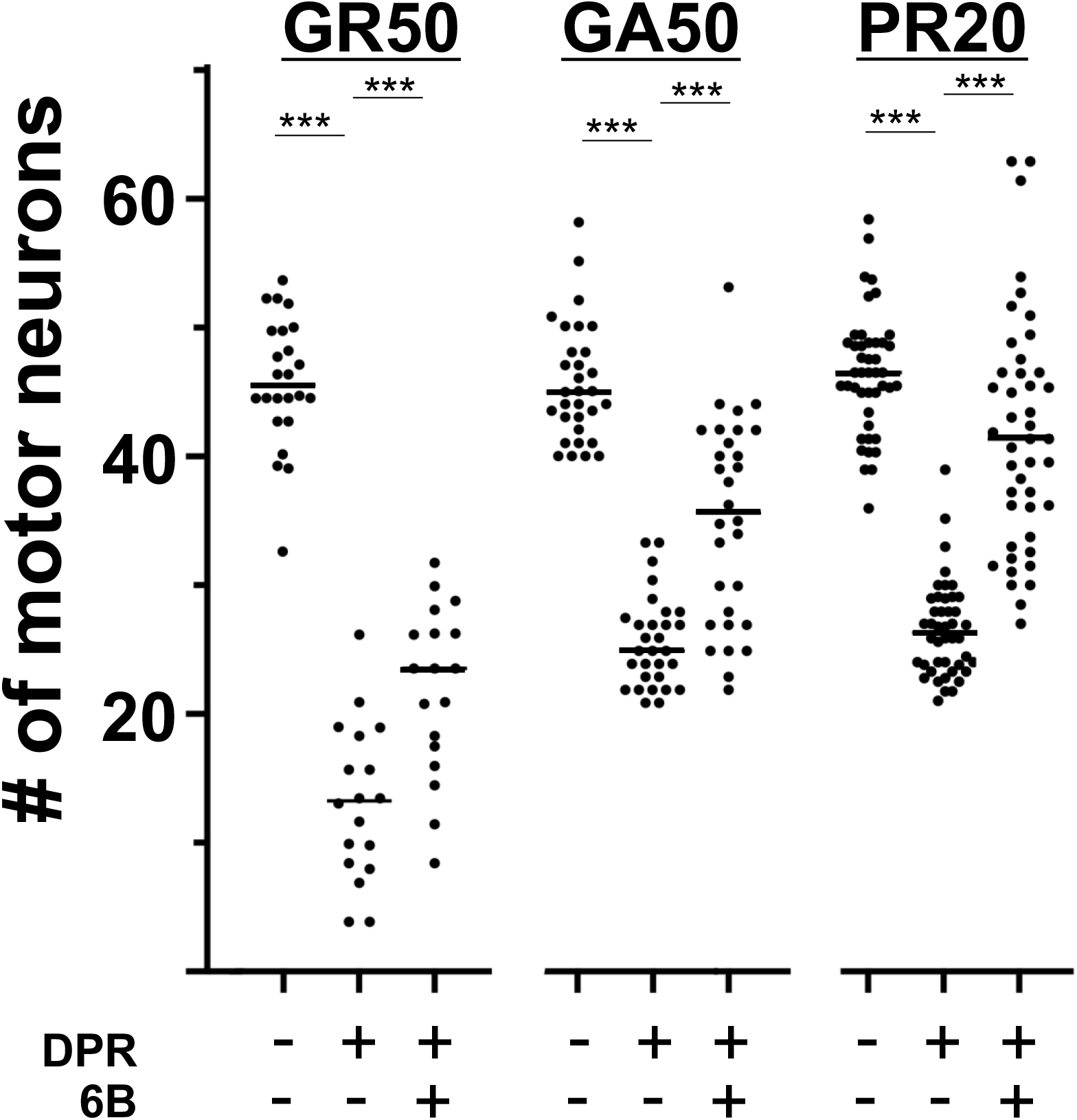
The SPOP inhibitor, compound 6B protects against DPR toxicity. DIV14 spinal cord neuron cultures were infected with HSV engineered to express the GA(50), GR(50) or LacZ or exposed to bath applied synthetic PR20 and were treated with either 6B (10 uM) or vehicle once. Five days later, cultures were processed for immunocytochemistry and motor neuron counts were obtained. By one-way ANOVA, DPRs lead to a statistically significant reduction in motor neuron number (compared with LacZ expressing cultures) and application of 6B leads to a statistically significant protection against DPR toxicity (e.g., GA(50) group, F(_2,69_) = 49.85, p < 0.0001; GR(50) group, F(_2,51_) = 114.0, p < 0.0001; PR20 group, F(_2,129_) = 79.32, p < 0.0001).

